# Robust estimation of recent effective population size from number of independent origins in soft sweeps

**DOI:** 10.1101/472266

**Authors:** Bhavin S. Khatri, Austin Burt

## Abstract

Estimating recent effective population size is of great importance in characterising and predicting the evolution of natural populations. Methods based on nucleotide diversity may underestimate current day effective population sizes due to historical bottlenecks, whilst methods that reconstruct demographic history typically only detect long-term variations. However, soft selective sweeps, which leave a fingerprint of mutational history by recurrent mutations on independent haplotype backgrounds, holds promise of an estimate more representative of recent population history. Here we present a simple and robust method of estimation based only on knowledge of the number of independent recurrent origins and the current frequency of the beneficial allele in a population sample, independent of the strength of selection and age of the mutation. Using a forward time theoretical framework, we show the mean number of origins is a function of *θ* = 2*Nμ* and current allele frequency, through a simple equation, and the distribution is approximately Poisson. This estimate is robust to whether mutants pre-existed before selection arose, and is equally accurate for diploid populations with incomplete dominance. For fast (e.g., seasonal) demographic changes compared to time scale for fixation of the mutant allele, and for moderate peak-to-trough ratios, we show our constant population size estimate can be used to bound the maximum and minimum population size. Applied to the Vgsc gene of *Anopheles gambiae*, we estimate an effective population size of roughly 6 × 10^7^, and including seasonal demographic oscillations, a minimum effective population size greater than 6 × 10^6^ and a maximum less than 3 × 10^9^.

## INTRODUCTION

Studying the differences in sequences between individuals in a population has the potential to give new insight into evolutionary processes: the evolutionary forces of selection, mutation, migration and drift can leave a signature in the pattern and frequency of polymorphisms in time and space, which population genetic theory can be used to infer [2, 8, 10, 11, 15, 22, 30]. A key parameter to estimate for any evolving population is the effective population size [9, 29], as it determines the underlying nature of the evolutionary dynamics and the relative importance of genetic drift versus selection for evolving traits. In particular, having an accurate estimate of recent effective population size has impact on our ability to predict the outcomes of evolution, as the current population size controls the mutational input through the parameter *θ* = 2*Nμ* and the fate of rare variants in a population via the population scaled strength of selection 2*Ns* [16]. However, there is not a single well-defined measure of effective population size and different estimates will depend on the particular evolutionary pressures on the trait or genomic region under consideration, as well as on previous population histories [5]. A common method to estimate effective population size is from the nucleotide diversity *π* of neutral regions of a genome, where for 2*Nμ* ≪ 1, we expect *π* < 2*Nμ* [5]. This relation represents a balance between mutations introducing variation at rate *μ* and drift removing variation at rate 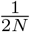. However, nucleotide diversity will tend to be dominated by population bottlenecks, and so be insensitive to recent population expansions [13], and there is a need for methods to estimate effective population sizes which are more representative of current day census size. Methods based on linkage disequilibrium tend to be limited to small population sizes [26]. On the other hand, although there are a number of methods that attempt to directly infer demographic history [4, 10, 11, 23], these methods are either complex and computationally intensive, or only able to detect long-term changes in population size. There are currently no methods that simply and robustly allow estimation of very recent effective population sizes.

A recent popular paradigm to study variation in populations are “soft sweeps”, where for sufficiently large population sizes (*N_μ_* ≳ 1) multiple copies of the same mutation, distinguished by their haploytpe background, co-exist in the population. This provides a direct genetic fingerprint on the rate at which mutations enter a population *θ*, which without their distinguishing haplotype backgrounds would be hidden. Precise information about *θ* is effectively hidden when mutations arise infrequently per generation (*Nμ* ≪ 1), since in this weak successive mutations regime, a single dominant haplotype fixes in a population before other haplotypes have a chance to establish; these are termed “hard sweeps”, as each subsequent sweep erases any previous information, giving a weak bound that *θ* ≪ 1. In a series of seminal papers by Pennings and Hermisson [12, 20, 21], much of the basic theory of soft sweeps was developed within a coalescence framework. In particular, the mean number and the distribution of independent origins in a neutral population sample were found to be given by Ewens’ sampling framework [7]. Estimating *N* from soft sweeps should be representative of the effective size over the time period of the sweep [13]. However, estimating the maximum likelihood effective population size requires using Ewens’ formula [7] for the probability of observing a certain number of distinct alleles in a sample of only neutral alleles, which although exact is not very practical for large sample sizes, as it requires evaluating the Stirling number of the first kind, a combinatorial factor that has not been implemented in most programming languages. In addition, when the mutant allele has not yet gone to fixation, we need to account for the fact that samples will contain both wild type and mutant alleles; this requires the extra complication of having to convolve Ewens’ formula with a binomial distribution for the probability of observing a given number of mutants in a sample given the frequency of the mutant.

In this paper, we present a simple semi-deterministic forward time approach, based on a non-homogeneous Poisson establishment rate of independent mutants, which thereafter grow deterministically [18]. We show that this gives very accurate estimates of the number of independent origins as a function of the time since selection sets in. In the haploid case we show explicitly the likelihood function is independent of the selection coefficient and only dependent on the frequency of the mutant allele, and so does not require estimation of the selection coefficient or the age of the allele. This approach has the advantage of being simple to implement, as the likelihood function is a non-homogeneous Poisson process, and is particularly appealing as the results can be understood in intuitive terms in a forward-time framework. Further, we show the method is robust to whether or not the mutation was pre-existing in the population, and is equally accurate for diploid populations with incomplete dominance (0 < *h* < 1). Finally, we apply our method to recent data from the Vgsc locus from the *Anopheles gambiae* 1000 genomes (1000Ag) project [1] to find an estimate of effective population size almost 2 orders of magnitude greater than is estimated by analysing nucleotide diversity. Moreover, to account for the marked seasonal population dynamics of this species, we show that it is possible to calculate a bound for the maximum and minimum effective population sizes, based on an estimate of effective population size using the constant population size method.

## THEORY

We calculate the likelihood of the number of origins with two assumptions: 1) we assume a non-homogeneous (time-dependent) Poisson process such that mutant alleles establish with rate *α*(*t*) = 2*N_μs_*(1 – *x*(*t*)), where *x*(*t*) is the frequency of all mutant alleles in the population; 2) after establishment of the *k^th^* mutant allele, its frequency *x_k_*(*t*) increases deterministically. The mean number of origins at time *T* is then determined by calculating the average number of establishment events in a time window 0 to *t_K_*, where *t_K_* is the latest possible time of establishment, such that it can grow deterministically to a critical frequency to be sampled from the population at some time *T*.

### Deterministic growth

We assume that the overall mutant population grows according to the following differential equation:

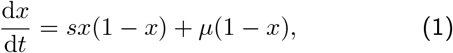

where the first term is the change in frequency due to frequency independent selection (assuming *s* ≪ 1) and second is the change in frequency due to mutations arising from the wild type population at mutation rate *μ*. This has the following closed form solution

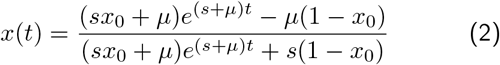

which in its tanh form is:

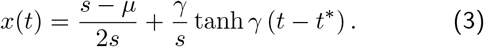

where

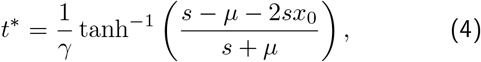

where *γ* = (*s* + *μ*)/2 and *x*_0_ is the initial frequency of the total mutant population. As in this deterministic framework the mutant allele only asymptotically reaches fixation as *t* → to, we identify *t*\* as the characteristic or typical time to fixation, which is the inflexion point of the tanh function and roughly the point at which the mutant has reached a frequency of (*s* – *μ*)/2*s* ≈ 1/2 for *s* ≫ *μ*; the actual time to fixation with discrete populations and drift will always be of the same order of magnitude as *t*\*. Here we assume that the initial frequency of the mutant is zero and so using the identity 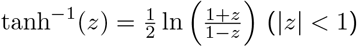,

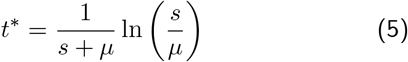

We see that the typical time to fixation *t*\* has a logarithmic dependence on the mutation rate, and can increase without bound for small mutation rates since we must wait for mutations to arise before selection can act to increase its frequency. Note that our approach here is in contrast to [13, 18, 27] who typically assume an expression for the mutant frequency which ignores initial conditions and de novo mutation, which as we see can cause a large effect on the time to fixation; in our case this is important as we require the mutant to have zero initial frequency, when the selection pressure arises.

### Stochastic establishment and likelihood of number of origins

We assume mutant alleles arise by de novo mutation at a time-varying (non-homogeneous) rate proportional to the number of wild type individuals *N_μ_*(1 – *x*(*t*)). De novo mutants must reach a critical frequency 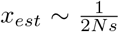 at which point more mutant individuals are added by selection compared to the change in number due to drift [6]. The probability that a de novo mutant, starting at frequency 1/*N*, grows by drift to size 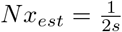, is just the inverse of the size of this neutral sub-population, *p_est_* ≈ 2*s*. The rate of establishment of mutants is then

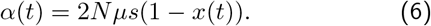

We make the assumption that establishments occur randomly and independently and so the underlying probability distribution for the number of establishments up to time *t_K_*(*T*), the time of establishment of last mutant to possibly be sampled at a latter time *T*, is given by a non-homogeneous Poisson process:

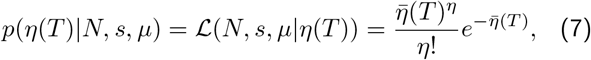

where *η*(*T*) is the number of independent origins at time *T*, and where the mean is given by the integral of the rate *α* up to time *t_K_*(*T*):

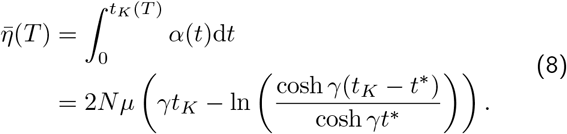

The time of the last establishment *t_K_*(*T*) is straightforward to calculate as shown next.

#### Calculating t_K_

The time for the last possible establishment, *t_K_* of the *K^th^* mutant, in order to be sampled with high probability at time *T*, is calculated by using a deterministic approximation for the change in frequency of the *K^th^* mutant. In an experiment, and in simulation, individuals of a population are sampled with a sample size *n_s_*; in simulation this is done using multinomial sampling with the allele frequencies determined from simulation. Here for simplicity we assume that when a mutant allele frequency is above *x_s_* = 1/*n_s_* then the mutant will be found in a sample of size *n_s_*. With a deterministic time-course of the *K^th^* mutant, there is a one-to-one correspondence between its frequency at time *T*, *x_K_*(*T*) and the time of establishment *t_K_*, given that its frequency must be *x_K_*(*t_K_*) = 1/2*Ns*.

To calculate *x_K_*(*t*), we use the fact that in the deterministic limit the ratio of the frequency of any mutant allele is fixed with respect to the overall mutant population, i.e. *x_K_*(*t*)/*x*(*t*) = *const*; this is true whenever the growth function of each mutant is of the same form 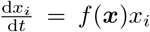, which can be proved by showing 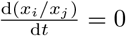. In this case, once a mutant arises in the population, we assume no more mutations can create the mutant from wild type and that there are no back mutations, so the growth of each mutant follows:

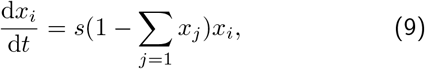

whilst the growth of the total number of mutants is given by Eqn.1; however, once the overall mutant population has established the effect of mutations will be weak compared to selection, as long as *s* ≫ *μ*, and so to a good approximation, the total mutant population also follows the same form as Eqn.9.

It is then simple to show that the frequency of the *K^th^* mutant is just a scaling of the frequency of the total mutant population *x*(*t*):

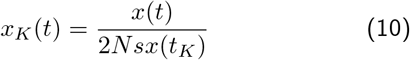

where we have used the fact that at the establishment time *t_K_* we know that the frequency of the mutant must be *x_K_*(*t_K_*) = 1/2*Ns*, and that *x_K_*(*t*)/*x*(*t*) = *x_K_*(*t_K_*)/*x*(*t_K_*). We then solve *x_K_*(*T*) = *x_s_*, for *t_K_* to give

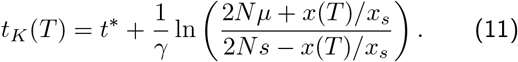

where we have again used the identity 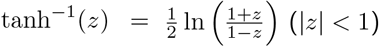 to arrive at this expression.

#### Simple expression for mean number of origins

The mean number of origins is calculated by inserting Eqn.11 into Eqn.8 and then after some algebra we find:

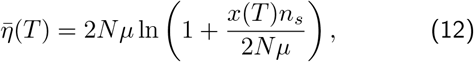

which we see only has dependence on the selection coefficient *s* through the frequency of the total mutant population *x*(*T*) at time *T*. This is consistent with the results in [20], where in the coalescence framework they find the probability of a soft-sweep in a sample size of 2, at fixation, is independent of the frequency sample path of the mutant allele and weakly bounded by selection through the fixation time. This result suggests that larger sample sizes *n_s_* increase the number of independent origins we should expect to observe.

As shown in the Supplementary Information, the theory can be extended to the diploid case, where we find an expression for the mean number of origins as a function of the dominance coefficient *h* (assuming incomplete dominance 0 < *h* < 1) and the selection coefficient *s*, as well as *N* and *μ*. In this case it is not clear whether the mean number of origins, and hence the Poisson distribution, is independent of the selection parameters *s* and *h*, as the resulting expression is complex. However, as we will see, the haploid expression is as accurate in the estimation of the effective population size as using the diploid expression, which suggests the dependence on *s* and *h* are weak. In addition, as shown by Pennings and Hermisson [20], the probability of a soft sweep has a weak ~ *s*^2^ dependence in diploid populations, which would also suggest a weak dependence on *s* for the number of origins.

## SIMULATIONS

### Methods

We simulate the population genetics of multiple recurrent mutations at a single locus using an infinite alleles Wright-Fisher process. Simulations start assuming a fixed wild type, so that the mutant frequency *x*(*t* = 0) = 0; each subsequent mutation that arises is given its own “allelic” identity to represent it arising on a different haplotype background, and once it enters the population the same allele cannot be produced by mutation from the wild type or any other allele. As is commonly assumed for an infinite-alleles process, we assume in addition there are no back mutations to the wild type. Each mutant allele has the same selective advantage *s* relative to the wild type. For population sizes up to *N* = 10^6^, we use multinomial sampling of alleles with fixed population size *N* to calculate the stochastic change in frequency between generations due to selection and drift. This is replaced by the equivalent multivariate Gaussian distribution with covariance matrix 〈Δ*x_i_*Δ*x_j_*} – 〈Δ*x_i_*〉〈Δ*x_j_*〉 = *x_i_*(*δ_ij_* – *x_j_*) for population sizes larger than 10^6^. Correspondence between the two methods was checked for simulations at smaller population sizes (not shown). In both cases mutations are treated separately and introduced with a non-homogeneous Poisson process, where the mean number of new mutant alleles in generation *t* + 1 is given by *N_μ_*(1 – *x*(*t*)), where *x*(*t*) is the frequency of all mutants in generation *t*; each of these new mutant alleles arise in the population with frequency 1/*N* (or 1/2*N* in the diploid case).

At various time points *T* we sample the vector of frequencies of all independent mutants *x*(*T*) = [*x*_1_(*T*),*x*_2_(*T*),*x*_3_(*T*),…, *x_K_*(*T*)], using multinomial sampling with *K* + 1 categories (including the wild type, which has frequency 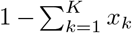), and sample size *n_s_*. This produces a sample vector *n*(*T*), where *n_k_*(*T*) is the number of the *k^th^* mutant sampled. The number of origins *η*(*T*) is then the number of different mutants that are non-zero in the sample.

### Results

In Fig.1 is plotted the time series of the frequency of each recurrent mutation from the Wright-Fisher simulations for *N* = 10^6^ and *s* = 0.05 and two different mutation rates, corresponding to 2*N_μ_* = 1 (A) and 2*Nμ* = 10 (B). We see that at the larger mutation rate there are correspondingly many more mutants in the population, and that the rate of production of mutants is proportional to the frequency of the wild type, signified by the lack of new mutants once the total mutant population has fixed. The red curve is a plot of Eqn.3, the deterministic solution for the total mutant population over time, and we see that it matches well the time-course found in the simulations, particularly for 2*N_μ_* = 10, where stochastic effects of the de novo generation of mutants becomes negligible. The frequency of each of the recurrent mutants follows the same scaling as the total frequency of all mutants, as assumed in the Theory section, and once the mutant population fixes, each of the recurrent mutants plateaus and stops changing in frequency (up to small relative fluctuations), which is as predicted by Eqn.9. In other words, in the deterministic limit there is a “crowding-out” effect, characteristic of logistic growth, where the growth of a mutant is limited by all other mutants in the population.

**FIG. 1.**
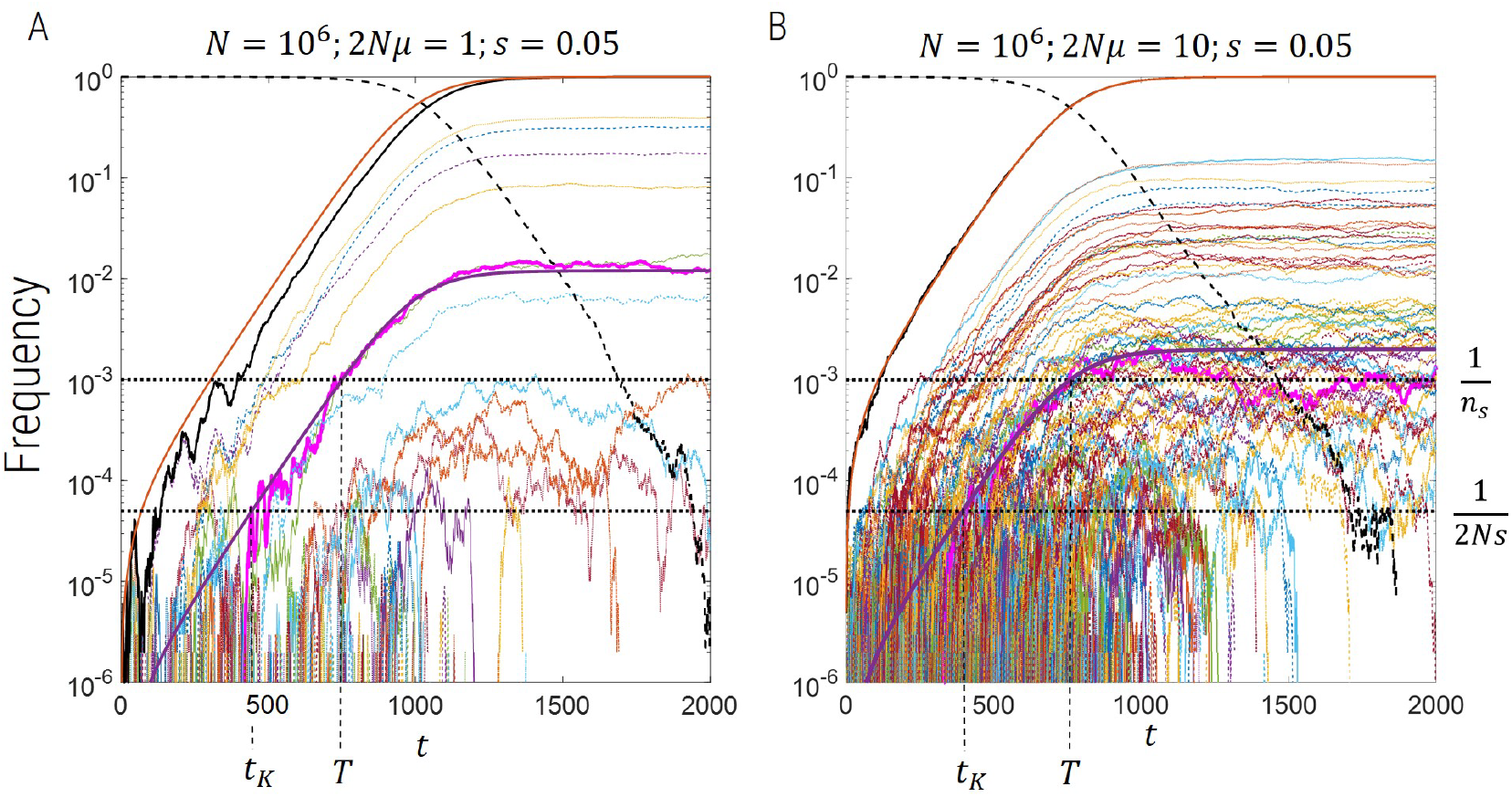
Time series of the frequency of each independent origin of the same recurrent mutant (range of different colours). A) *N* = 10^6^, 2*Nμ* = 1, & *s* = 0.05, B) same as A, but with 2*Nμ* = 10. Solid black line is the sum of all mutant frequencies 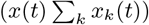, dashed black line the frequency of the wild type (1 – *x*(*t*)), and the solid red line is the deterministic time course given by Eqn.3.

In each plot the highlighted mutant in the thick magenta solid line shows an example of a mutant establishing at the frequency 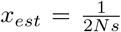, at time *t_K_*, and then reaching the critical sampling frequency at a time *T*. If *T* is the time of sampling, then this would be the last possible mutant that could contribute to a sample, and the time between 0 and *t_K_* would be the window over which mutants can be generated that could contribute to a sample at time *T*. The distribution of the number of origins at time *T* is just the distribution of the number of establishments in this time window; this is the basis of the semi-deterministic theoretical calculation of the number of origins described above.

Fig.2 shows the results for the mean number of origins 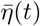 calculated from simulation (squares), compared the semi-deterministic theory presented in this paper (thick lines) and Pennings and Hermisson’s calculation [20] based on Ewens’ sampling theory [7]. We see in general that the time-course of 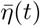 reflects the time-course of the frequency of the total mutant population, with a sigmoidal variation, where for the largest selection coefficients we see a plateau reached in less than 500 generations. Both the semi-deterministic theory and Ewens’ theory predict that the plateau of 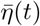 is independent of the selection coefficient, since 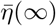 is roughly given by time window over which mutants can be generated, which approximately scales as 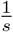, multiplied by the rate of establishment of mutants, which scales like ~ *s*, cancelling the *s* dependence. We see that the simulations agree with this prediction for the larger population sizes, but for *N* = 10^6^, the number of origins decreases for long times; this is due to drift removing very low frequency variants at the smaller population size, whilst at the larger population sizes drift acts more slowly, such that the change is insignificant on the timescale of the simulation. Finally, we see that the time-course of the mean number of origins before the plateau is different for each population size, where for the smaller selection coefficients the mean number of origins arise more slowly for larger population sizes. This is related to the deterministic time-course of the mutant frequency which, given the initial condition that the mutant frequency is zero, has a strong dependence on the mutation rate as shown by Eqn.5. The simulations are performed for fixed 2*Nμ*, and so a larger population size means a smaller mutation rate and so 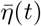 increases more slowly; however, at long times the plateau number of origins agree for all population sizes (not shown), as predicted by Eqn.12.

**FIG. 2.**
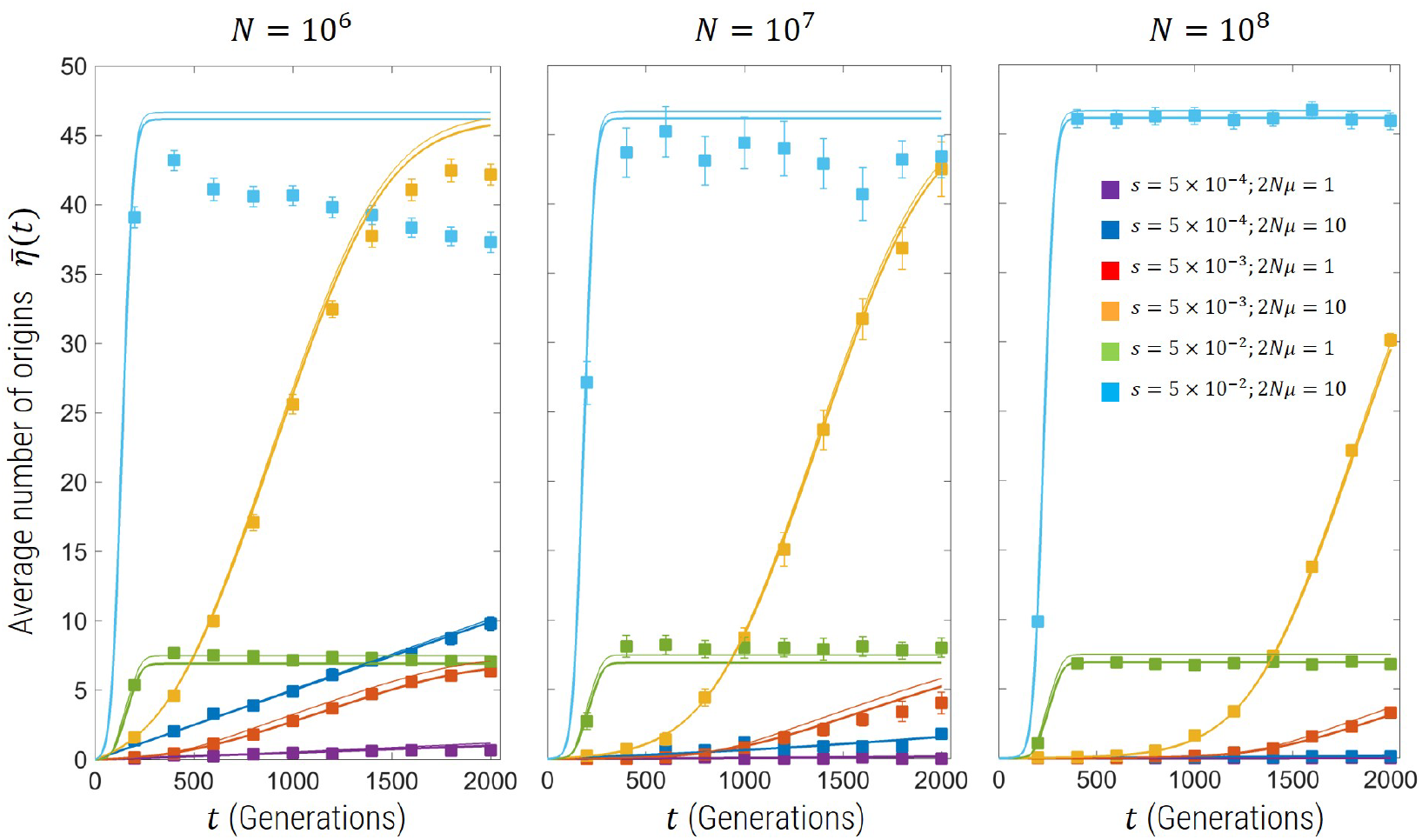
Average number of origins for population sizes of *N* = 10^6^, *N* = 10^7^, and *N* = 10^8^. The squares show the simulation results and standard error bars for the parameter combinations shown in the legend; for *N* = 10^6^ and *N* = 10^7^ the simulations used multinomial sampling of the Wright-Fisher drift process with 50 and 10 replicates, respectively, for each parameter combination, while for *N* = 10^8^ the multinomial sampling is replaced by the multivariate Gaussian distribution approximation of the drift process (see subsection Methods above), where 100 replicates are used in this plot. The solid thick lines are the predictions for the same parameter combination of the semi-deterministic theory described in this paper (Methods), while the thin lines represent the prediction of Pennings and Hermisson [20], based on Ewens’ sampling theory [7].

We also examine the distribution of the number of origins in Fig.3 from Wright-Fisher simulations (1000 replicates) at a population size *N* = 10^8^, selection coefficient *s* = 0.05, and mutation rates 2*Nμ* = {0.1,1,10}. The theory presented in this paper describes the distribution very well for all times up to and including fixation. On the other hand Ewens’ sampling framework predicts in a sample of *n_s_* neutral alleles that the distribution of the number of distinct mutant alleles *η* is

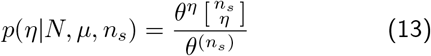

**FIG. 3.**
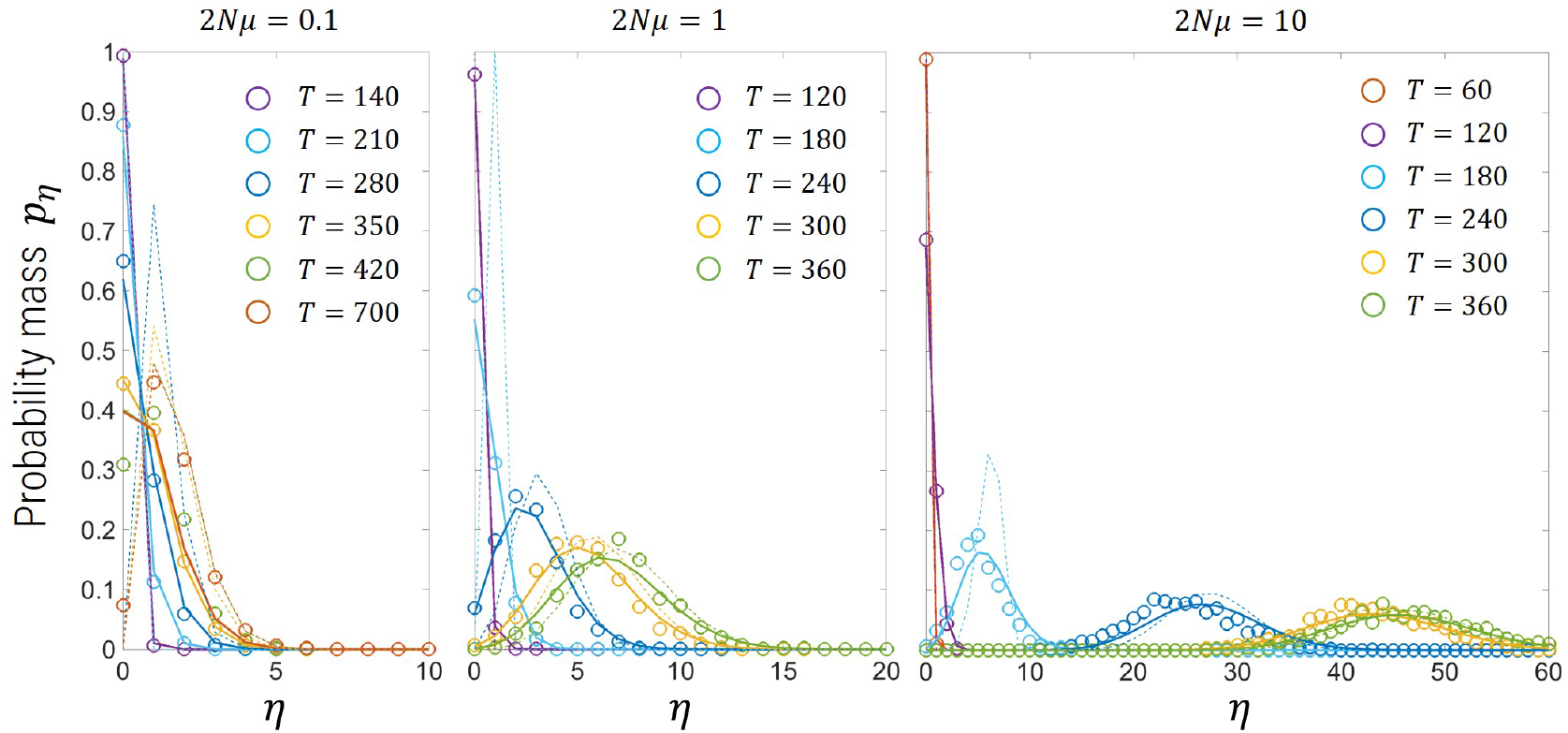
Distribution of the number of origins for simulations with various mutation rates for *N* = 10^8^ and *s* = 0.05 (open circles) compared to theory in the this paper Eqns.12 & S15 (solid lines) and Ewens’ sampling formula (dotted lines). For the mutation rates 2*N_μ_* = {0.1,1,10} the corresponding typical fixation time (Eqn.5) is *t*\* ≈ {370, 320, 280} generations.

where 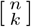 is the unsigned Stirling number of the first kind, which is a combinatorial factor which arises in the expansion of the rising factorial 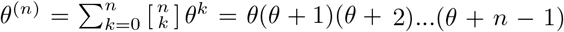. However, if the mutant allele has not fixed then the probability distribution of *η* mutants alleles is the convolution of Eqn13 with a binomial distribution that in a sample of size *n_s_* we see *n_x_* mutant alleles given a frequency *x*(*t*) of the mutant population. This convolution has no known closed-form solution and for large sample sizes is computationally intensive. In Fig.3 the dotted lines are a plot of Ewens’ theory Eqn.13 without this convolution and *n_s_* replaced in Eqn.13 by *n_s_x*(*t*) (calculated in Mathematica [28]) and as expected it does poorly when the mutant hasn’t yet fixed, and is quite accurate at later times when the mutant is near or at fixation. When the mutant allele is at fixation, the semi-deterministic likelihood of this paper and that from Ewens’ formula are closely matched (Fig.6).

## PARAMETER ESTIMATION

### Haploid

As described above, the semi-deterministic theory calculates the likelihood function for the number of observed independent origins, and we find it is only a function of 2*Nμ*, the frequency of the mutant population at the time of sampling *x*(*T*) and the sample size *n_s_* = 1/*x_s_*. Typically, the mutation rate will have been independently determined, and so we can determine a maximum likelihood estimate of *N* given knowledge of the *n_s_* and *x*(*T*), which can be estimated from the sample. In Fig.4A is the log_10_-error of this estimation process using 100 replicate Wright-Fisher simulations, with sample size *n_s_* = 1000, where the true *N* is known. We see that for mutant frequencies *x* > 0.1, the error of our estimate *N*\* is always less than a factor of 10^0.2^ ≈ 1.6, which means the effective population size is accurately determined to much less than an order of magnitude. Moreover, the accuracy increases for increasing 2*Nμ*, where it is less than 10^01^ ≈ 1.3 for 2*Nμ* ≥ 10.

**FIG. 4.**
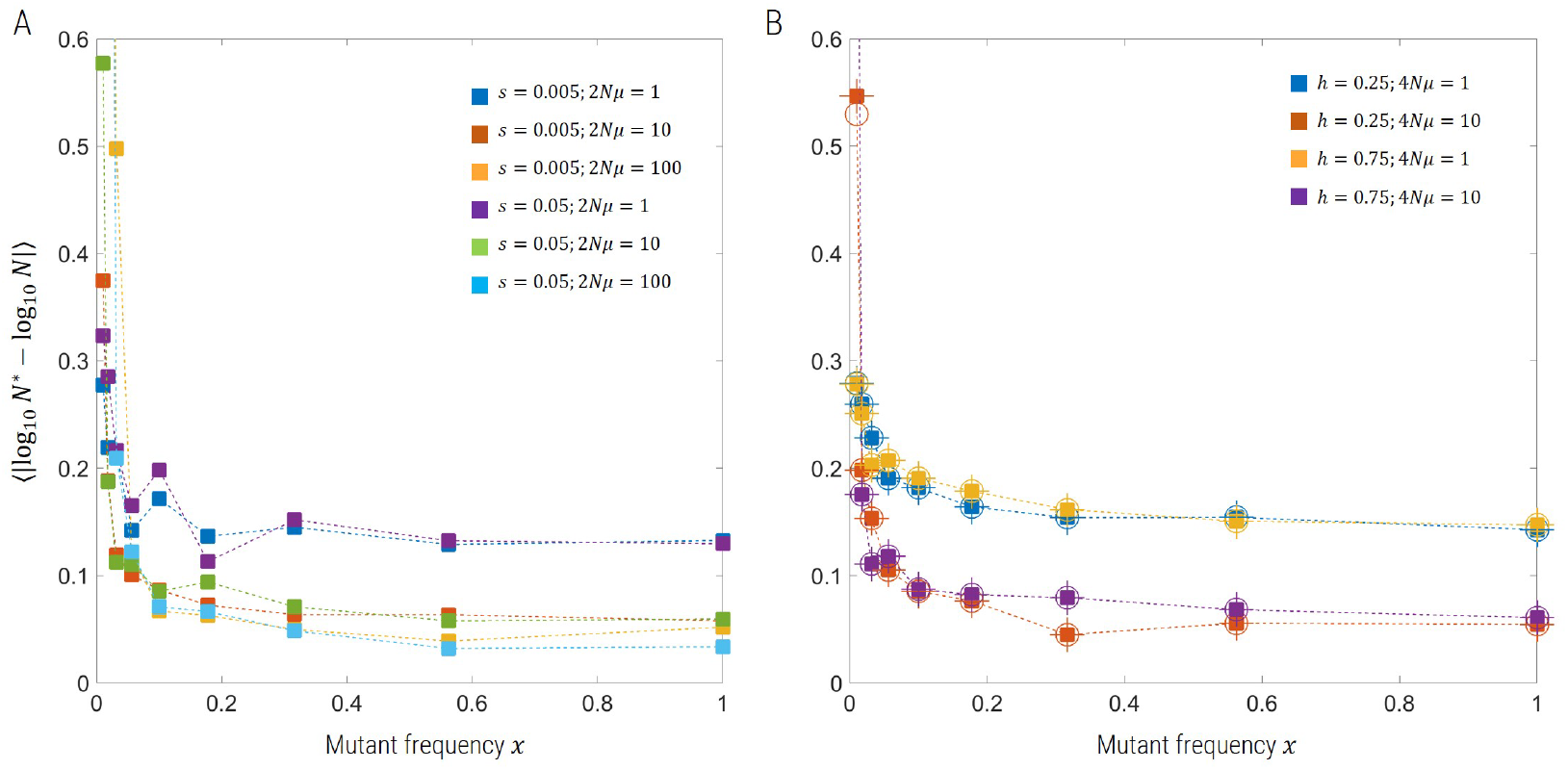
log_10_-error in estimating the true effective population size, for A) Haploid populations with *N* = 10^8^, B) Diploid populations with *N* = 5 × 10, for various selection coefficients, mutation rates, and dominance coefficients (diploid only) from Wright-Fisher simulations (100 replicates for each parameter combination). A) We use Eqns.12 & S15 to determine the maximum likelihood estimate. B) For the diploid population we use the same Poisson likelihood function, but with mean given by Eqn.13&14 in the Supplementary Information, where we assume perfect knowledge of *T* (squares) and also compare to the case where we have a systematic error in our knowledge of *T*, where the true time is *T*/2 instead *T* (circles), and we see the estimates are unchanged. In addition, for the diploid population we use the haploid likelihood function (Eqns.12 & S15) to estimate 2*N* (plus signs) and find again excellent agreement.

### Diploid

We can also accurately estimate the effective population size from diploid simulations. As described in the Supplementary Information, we extend the semi-deterministic theory to the diploid case with incomplete dominance (0 < *h* < 1) by using the exact implicit solution *t*(*x*) for how the frequency *x* of the mutant allele changes over time to calculate time of establishment of the last mutant to be sampled at some later time *T*. This is then used to calculate the likelihood function *p*(*η*|*N, s, h, μ*), where we assume a known mutation rate. We are still left with having to jointly estimate *N, s* and *h* in the diploid case. However, we expect that the dependence on *h* and *s* will be weak ([20]), although it is not straightforward to show this explicitly, as in the haploid case, where there is no dependence on *s*, even before fixation. To show this we use the implicit relation (Eqn. 2 Supplementary Information) to numerically estimate *s*\* that gives *t*(*x*) = *T*, where we assume perfect knowledge of the dominance coefficient *h*. We see in Fig.4B that the estimate of the effective population size from diploid simulations has a similar accuracy as the haploid simulations, and is robust to knowledge of the exact time selection sets in *T*; the error is taken up in the estimate of *s* (not shown). We also use the haploid semi-deterministic theory to estimate the effective population size, accounting for double the number of chromosomes, shown by plus signs in Fig.4B; again we see that the estimate of *N* is identical using the haploid method for a given set of parameters, *s, h* and *μ*. Both the robustness of estimates to the exact knowledge of *T* and that the haploid gives identical estimates indicates that the direct dependence on *s* and *h* is very weak or non-existent, at least for weak absolute selection [20].

### Haploid with pre-existing mutations

Finally, we examine the effect that pre-existing mutations have on our estimate of the effective population size. We run simulations such that for times *T_d_* < *t* < 0 the mutant allele has a negative selection coefficient *s* = −*s_d_*, where 2*Ns_d_* = {0,10^3^,10^4^,10^5^,10^6^}, *T_d_* = −1000 generations and *N* = 10^8^, *s* = 0.05 and 2*Nμ* = 1. The mean number of origins 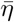 is plotted in Fig.5A, for the various values of *s_d_* as well as for the case of no pre-existing mutations (black hexagram symbols); we see that as the mutant allele becomes increasingly neutral before positive selection sets in, the number of origins is larger, except for long times where the plateau of 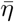 is approximately independent of *s_d_*. This suggests the overall effect of pre-existing mutations is to cause a time advance on the number of origins. This again would suggest that the estimate of effective population size should be robust to pre-existing mutations, which we see to be the case in Fig.5B, where the error in estimating *N* using Eqn.12 for the mean of the Poisson likelihood function is roughly independent of *s_d_* and very similar to assuming no pre-existing mutations (black hexagrams).

**FIG. 5.**
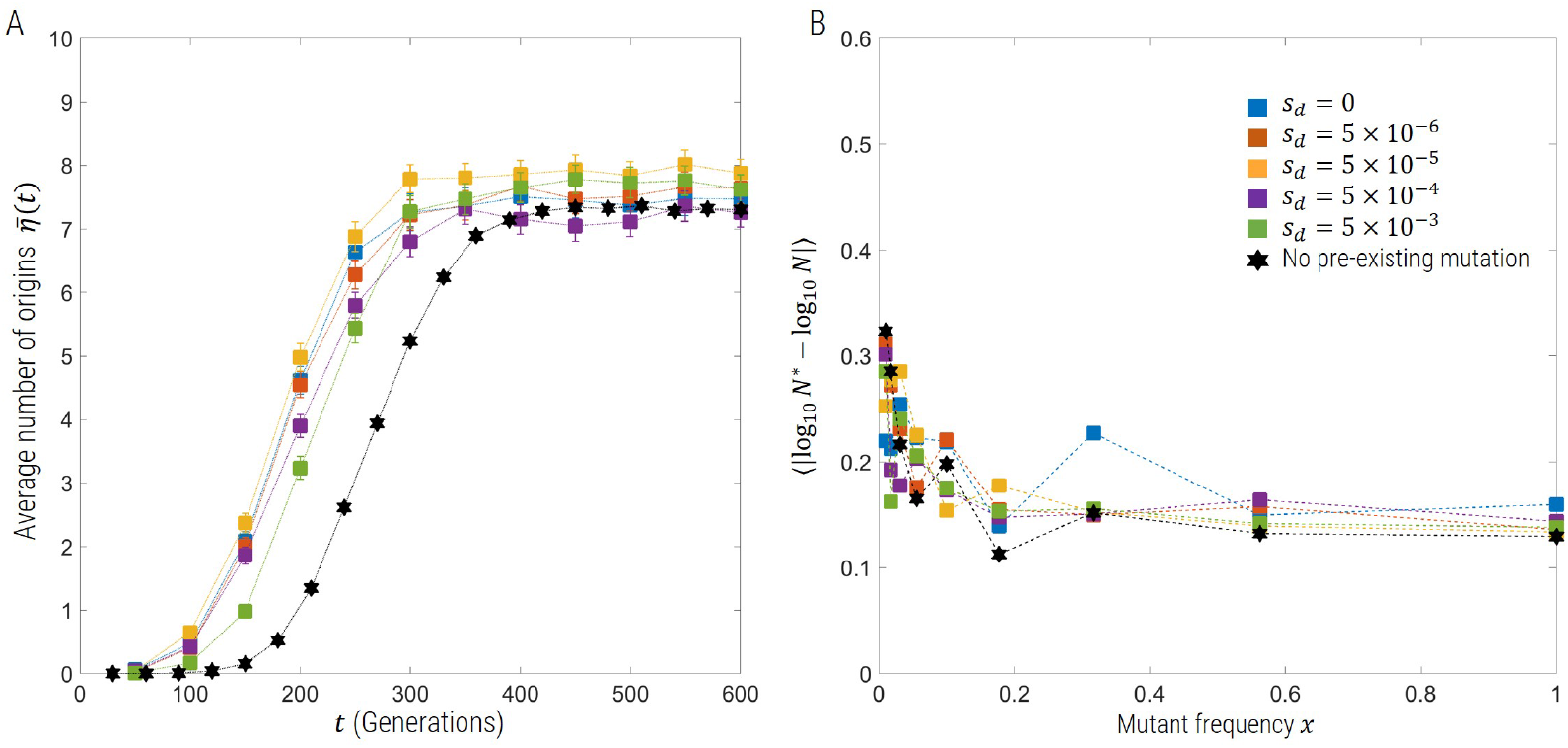
Mean number of origins for haploid simulations with pre-existing mutations (A), where the black hexagram symbols represent simulations without pre-existing simulations, and (B) log_10_-error in maximum likelihood estimate of the true effective population size *N* = 10^8^ from Wright-Fisher simulations with various values of the deleterious selection coefficient *s_d_* (100 replicates for each parameter combination).

### APPLICATION TO DATA FROM AG1000 PROJECT

Recently published data from the *Anopheles gambiae* 1000 genomes project (Ag1000) has extensive population level sampling of the genomes of mosquitoes across sub-Saharan Africa [1]. The gene for the voltage-gated sodium channel (Vgsc) is known to have at least two single nucleotide mutations in the same codon that confer resistance to insecticides, *L995S* (2984*T* > *C*) and *L995F* (2985A > *T*), and phylogenetic analysis of this gene reveal 10 haplotype clusters (Fig.4 in ref[1]) with a current mutant frequency of *x* ≈ 0.78 determined directly from the data. If we assume either mutation is required for resistance, this gives a mutation rate of *μ* ≈ 6 × 10^-9^, assuming a base-pair mutation rate of 3 × 10^-9^, which is based on a recent accurate estimate from *Drosophila* [14], as the mutation rate has not been directly measure for *Anopheles gambiae*. Applying the haploid algorithm to this data, accounting for the factor of 2 between chromosomes and individuals, and using a sample size of *n_s_* = 1530 chromosomes from 765 mosquitoes, gives an effective population size *N* = 6.2 × 10^7^ (2.7 × 10^7^, 1.2 × 10^8^), where the values in brackets are the 95% confidence intervals (2 ln units from max likelihood), as shown in the plot of the likelihood function in Fig.6. This estimate is almost 2 orders of magnitude greater than that of *N* ≈ 10^6^ from a nucleotide diversity *π* ~ 0.01. In the same paper, the authors use the more sophisticated “stairway” plot [10] and *∂a∂i* [11] method to estimate population history, and find most recent effective population sizes of order *N* ≈ 10^7^, which is roughly 6 times less than our estimate.

**FIG. 6.**
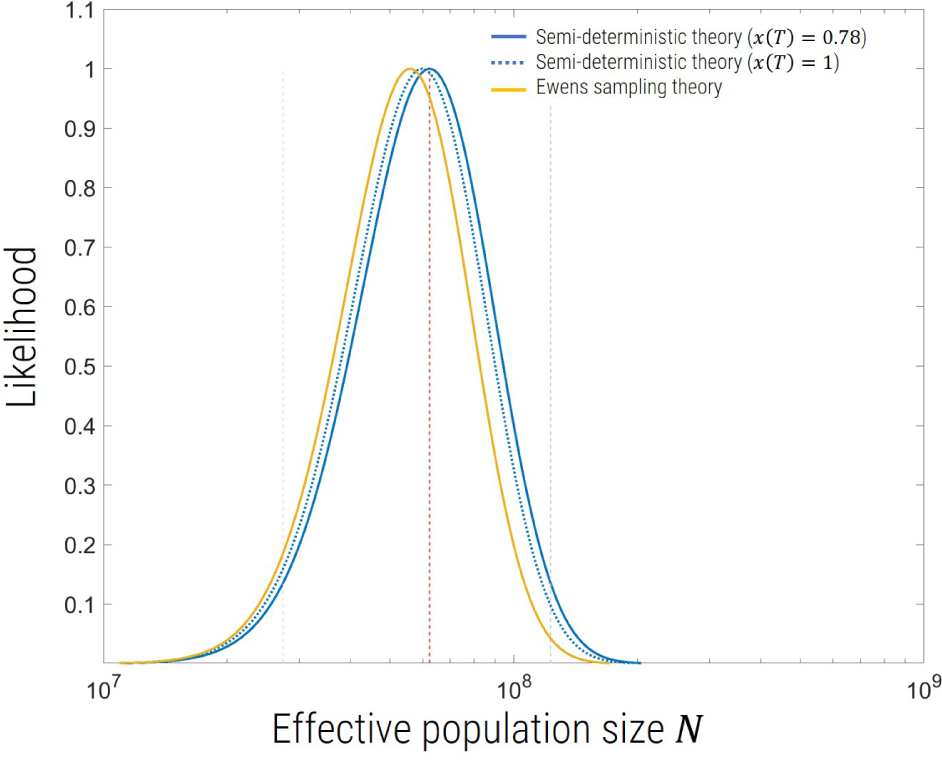
Likelihood (normalised) of the number of origins as function of effective population size given an observed number *η* = 10 and samples size *n_s_* = 1530 chromosomes, corresponding to that found for the Ag1000 project [1] for the Vgsc resistance locus. As shown in the legend, the semi-deterministic theory in this paper, assuming a current day frequency of *x* = 0.78 (as observed) is compared to assuming *x* = 1 and the Ewens’ sampling theory Eqn.13, which only has applicability for *x* = 1. The 95% confidence intervals (grey dotted lines) and maximum likelihood effective population size (red dotted line), are shown for the semi-deterministic likelihood function with *x* = 0.78.

Note that we can also apply the method to each resistance mutant separately *L995S* and *L995F*, which have frequencies of ≈ 0.28 and ≈ 0.5, and 5 independent origins each, which assuming a single base-pair mutation rate of ≈ 3× 10^-9^ for each of these, gives the following estimates of effective population size *N* = 6.6×10^7^ (1.9×10^7^,1.7×10^8^), and *N* = 6.0× 10^7^ (1.8× 10^7^,1.5× 10^8^), respectively, where the values in brackets, are again the 95% confidence intervals. We see the estimate based on each SNP are consistent with the estimate above based on both SNPs, but, as expected, with larger confidence intervals.

However, it is known that in many sub-Saharan regions mosquitoes undergo seasonal demographic changes, where the population size changes between wet and dry seasons by up to a factor of 100 [3, 17, 19, 25]. To check the impact of demographic changes on our population size estimates, we ran simulations for a mutant with *s* = 0.05, with an oscillating population size 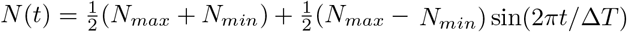, with a period of Δ*T* = 10 generations, which is approximately 1 year and much shorter than the expected time to fixation of the mutant of approximately 300 generations (Eqn.5). The simulations were performed with various peak-to-trough ratios *ϕ* = *N_max_*/*N_min_* = {10,100,1000} and with two constraints: 1) that the geometric mean 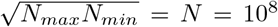 and 2) that the harmonic mean 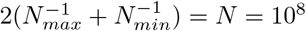. Simulations with constrained arithmetic mean were also performed but are not shown. We see in Fig.7, that for *ϕ* ≤ 1000, simulations that constrain the geometric mean gives fewer independent origins than with constant populations, and simulations that constrain the harmonic mean give more independent origins. This conversely means that for oscillating demographic changes the estimate *N*\* of the effective population size, using the constant population size theory of this paper, will be an *underestimate* of the geometric mean of the population and an *overestimate* of the harmonic mean. This allows us to derive simple expressions for bounds on the maximum and minimum effective population sizes given our estimate *N*\* and *ϕ*:

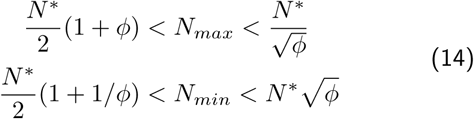

**FIG. 7.**
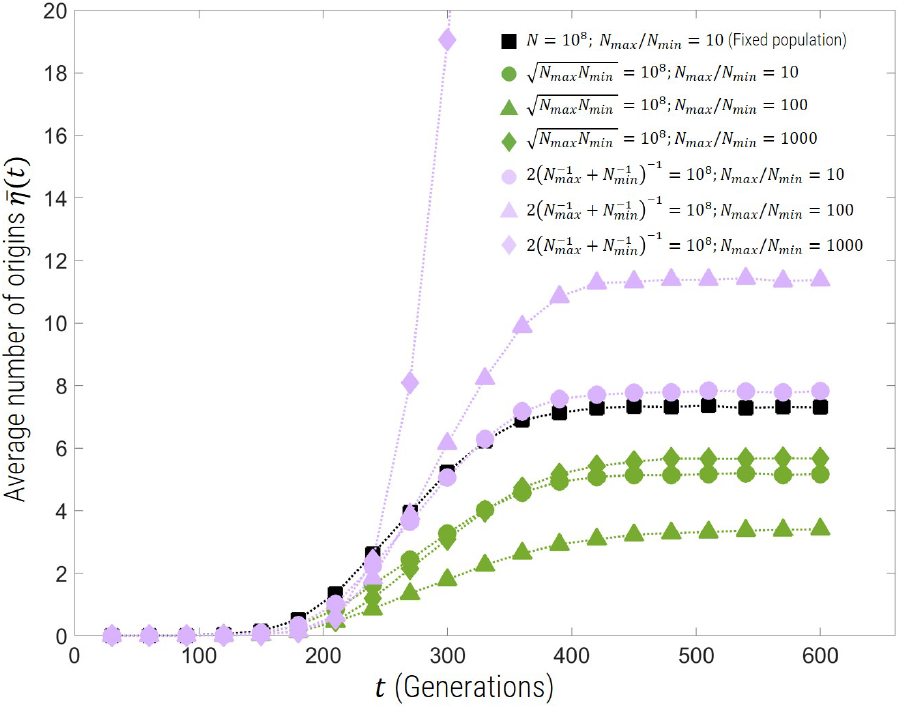
The mean number of origins from Wright-Fisher simulations (1000 replicates) for oscillating population size with period Δ*T* = 10 generations, selection coefficient *s* = 0.05, 2*Nμ* = 1 and with the geometric mean (green) and harmonic mean (purple) of *N_max_* and *N_min_* constrained to 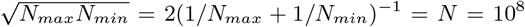, for different peak-to-trough ratios. Black squares represent constant population size simulations. We see that the constant population size simulations are bounded from above by simulations constrained to have the same harmonic mean, and bounded from below by the simulations with constrained geometric mean.

Using the estimate above *N*\* = 6.2 × 10^7^, we arrive at the following bounds for the maximum and minimum population sizes, assuming *ϕ* = 100: 6.2 × 10^8^ < *N_max_* < 3.1 × 10^9^ and 6.2 × 10^6^ < *N_min_* < 3.1 × 10^7^. So we see that our analysis estimates that the peak population size can’t be greater than 3.1 × 10^9^ and the minimum population can’t be less than 6.2 × 10^6^. Note that in Fig.7 the number of origins with constrained geometric mean varies non-monotonically as *ϕ* increases, so for very large *ϕ* we find the number of origins exceeds the constant population size scenario (not shown), and so this bound will not work. Simulations that constrain the arithmetic mean of the maximum and minimum population sizes show that the number of origins monotonically decreases with increasing *ϕ*, but are significantly less than even the constrained geometric mean case (not shown). This means our estimate *N*\* will be less than the arithmetic mean for all *ϕ*, but the equivalent to Eqn.14 provides a much weaker *lower* bound on *N_max_* and *N_min_*.

## DISCUSSION

Estimating the recent effective population size is of paramount importance to understanding and predicting the evolutionary dynamics of natural populations. As has been previously suggested [13], methods that estimate effective population size based on nucleotide diversity are likely to give estimates which are much smaller than the current day census size, as such metrics are dominated by historical population bottlenecks. Although methods based on linkage disequilibrium can detect recent effective population sizes, they tend to be limited to small populations [26]. In addition, methods that estimate demographic histories tend to be computationally complicated and with limited range of applicability, such as only detecting long-term variations [10, 11, 23] or limited to small population sizes [4]. However, a genomic region undergoing current selection should leave a signature which represents an effective population size more representative of the census size during the sweep [13]. When the mutational input into a population is large 2*N_μ_* > 1, we expect a signature of a selective sweep will be a large diversity of haplotype backgrounds, due to multiple and recurrent independent instances of the same mutation that is under positive selection; such a sweep has been termed a soft-sweep as multiple rather than a single haplotype dominate the sweep [12]. Although Pennings & Hermisson seminal work [20, 21] laid out much of our understanding of soft-sweeps within a coalescence framework, many quantities like the likelihood of the number of origins, particularly when the mutant population has not yet fixed, are not straightforward to calculate numerically.

In this paper we have presented a simple semi-deterministic haploid forward-time theory of the number of independent origins of a recurrent mutation. We show that the distribution of the number of origins is very closely approximated by a Poisson distribution with a mean number of origins that has an exact and simple closed-form solution for the haploid case, which is independent of the selection coefficient and the age of the allele, and only depends on 2*Nμ*, the sample size and the current day mutant frequency. We show it works robustly for diploid populations with incomplete dominance, and whether or not mutations are preexisting in the population before the selection pressure arose.

Our forward-time semi-deterministic theory also provides an intuitive insight into the dynamics of soft sweeps, where it is clear there is a demarcation between the stochastic and deterministic stages for each haplotype contributing to a soft sweep. New origins are generated by recurrent mutation, and these must establish by growing to a frequency where deterministic selection outweighs drift; thereafter growth is approximately deterministic of each independent mutant. The deterministic part of the theory shows that at sufficiently large population sizes the growth of each recurrent mutant is just a scaling of the overall mutant population and grows logistically, where other mutants “crowd-out” the growth of a particular mutant; once the wild type is extinct new mutants cannot arise, and growth of each recurrent mutant is zero, so this structure is effectively frozen, which is confirmed by simulation up to small fluctuations due to drift. Including drift in this picture means that this frozen structure is only temporary as drift will take of order *N* generations to act. This is seen in the simulations at even a moderate population size of *N* = 10^6^, where drift can act on the small frequency variants causing a decrease in independent origins for long times; however, for very large populations *N* ≫ 10^7^ there is a stable plateau as predicted by the theory. This suggests that Ewens’ sampling theory and the calculation in this paper will not be valid for small populations after fixation of the mutant, since the supply of mutants has been switched off; therefore the semi-deterministic approach in this paper will be limited to times at or before fixation for small population sizes.

The framework of this semi-deterministic theory also makes clear why selection should have little effect on the plateau number of origins, as the rate of establishment is proportional to the *s*, whilst the time window over which new origins can be generated is proportional to the lifetime of the wild type, which scales as 1/*s*, giving a number of origins that is independent of *s*. In addition, our result for the mean number of origins shows further that it is only dependent on the selection coefficient through the frequency of the mutant population, and in particular on the ratio of the number of mutants in the sample to the number of new mutants that enter every generation (2*Nμ*). Surprisingly, as found by Pennings & Hermisson [20] the number of origins does not depend on the exact sample path (history of the population frequency) of the mutant; here we see further that the number of origins only depends on the frequency of the mutant at a given time.

Finally, we estimated the effective population size of *Anopheles gambiae* and *Anopheles coluzzii* to be approximately *N* ≈ 6.2 × 10^7^ using data from the 1000Ag project [1], which is roughly 2 orders of magnitude larger than estimated using the same underlying data from nucleotide diversity and much closer to what is likely to be the census population size in recent history. This supports simple calculations of Karasov et al. [13], which suggested values of effective population size derived from nucleotide diversity are too small to explain adaptation of resistance alleles or the occurrence of multiple resistance haplotypes for the *Ace* gene in *Drosophila melanogaster*. Here, we have provided a very simple and robust method to quantify this effect. The demographic history of *Anopheles* has also been estimated from the 1000Ag project data [1] using the “stairway” plot [10] and *∂a∂i* [11] methods, giving a recent population size of roughly *N* ≈ 10^7^, greater than the nucleotide diversity estimate, but smaller than our estimate. A possible reason for this discrepancy is that these methods tend to detect long-term demographic changes, so that the difference could represent recent population growth in the past 100 years. However, the estimates in [1] are based on applying each of these methods to data from each geographic region, whereas the estimate here is based on data from all geographic regions in the Ag1000 data. In the completely panmictic case, the estimate in each region should agree with the estimate based on pooling the data, but as discussed below if there is spatial structure then the relation between the two estimates would not be straightforward.

It is also known that the Anopheles populations undergo seasonal demographic fluctuations with peak-to-trough population sizes of order 10 – 100 [3, 17, 19, 25].

To investigate the effect of such fluctuations on our population size estimates we performed simulations of oscillating population sizes over time for peak-to-trough factors less than 1000. These results showed that a constant population size estimate will tend to underestimate the geometric mean of the maximum and minimum of the population size, whilst overestimating the harmonic mean, allowing a quantification of bounds on the maximum and minimum population size of 6.2 × 10^8^ < *N_max_* < 3.1 × 10^9^ and 6.2 × 10^6^ < *N_min_* < 3.1 × 10^7^, assuming a peak-to-trough ratio of 100.

The results of these oscillating demographic simulations are in contrast to those of Wilson et. al. [27], which showed that the probability of a soft sweep in a sample size of 2 only depends on the cycle-averaged harmonic mean, when demographic oscillations are fast. It is simple to show that the cycle-averaged harmonic mean is just the geometric mean of the maximum and minimum population sizes; however, our results show different peak-to-trough ratios give significantly different numbers of independent origins for the same geometric mean. This suggests the probability of a soft-sweep in a sample size of 2 is a weak measure of the diversity of haplotypes compared to the number of independent origins.

Our estimation also makes the assumption that the populations are well-mixed or panmictic and constant over time, which clearly requires testing regarding the Ag1000 data, which consists of the sequences of individuals collected over the wide spatial region of sub-Saharan Africa. As discussed by Ralph & Coop [24], we would expect our results to be accurate in the limit of strong long-range or non-local dispersal, which mimics the panmictic approximation; on the other hand if local migration is strong, spatial structure of the populations would tend to give a larger number of origins compared to the panmictic case, which would suggest our method would overestimate the effective population size needed to explain an observed number of origins. In other words, it is possible that spatial structure could account partially or wholly for the large number of origins observed in natural populations of *Anopheles gambiae* and *Anopheles coluzzii*. Further theory and simulations will be needed to test this hypothesis.

## I. ACKNOWLEDGMENTS

We thank Alistair Miles for help with analysing the 1000Ag data and useful discussion including Tin-Yu Hui and Nicholas Harding. This work was supported by grants from the Bill & Melinda Gates Foundation and the Open Philanthropy Project.

## Supplementary Information

### MEAN NUMBER OF ORIGINS FOR DIPLOID POPULATION

In the diploid case an exact analytical form for the total mutant frequency *x*(*t*) does not exist, however, we can make progress by using the implicit solution *t*(*x*), which does have an exact analytical form. The differential equation determining the change in frequency of the mutant population, including diploidy is:

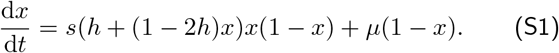

Using partial fractions the implicit solution is:

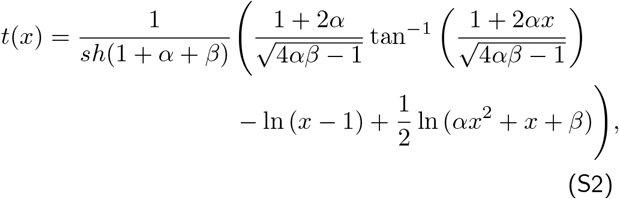

where 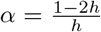 and 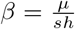. Using a similar argument as the haploid case, we can find the frequency of the last or *K^th^* mutant (equivalent of Eqn.10 in main text):

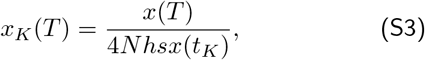

where here we have used the approximation that the establishment frequency will be dominated by heterozygotes, and be 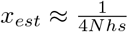. Eqn.S2 is then used to solve for *t_K_*, using as before *x_K_* (*T*) = *x_s_*:

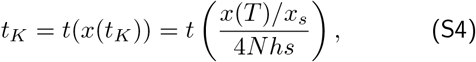

where *t*(*x*) is the implicit function given in Eqn.S2. Similarly, we approximate the probability of establishment as that of the heterozygote, so *p_est_* ≈ 2*hs*; this gives the in-homogeneous Poisson rate as

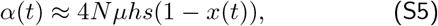

and so the mean number of origins will be

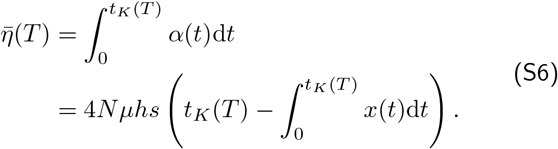

However, we do not have an explicit form for *x*(*t*). Given the implicit form (Eqn.S2) this can be numerically integrated, however, we instead develop an approximation for *x*(*t*). This involves approximating the RHS of Eqn.S2, *F*(*x*) by a piecewise quadratic, up to a frequency *x*^*^, which is the frequency at which 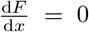; for 0 ≤ *h* ≤ 1 this is a reasonable approximation. As this approximate *F*(*x*) is quadratic, an exact solution can be found for each region, 0 ≤ *x* ≤ *x*^*^ and *x*^*^ ≤ *x* ≤ 1, and is of the form given for the haploid case in Eqn.3 in the main text. These solutions are then matched at the common point of inflexion which occurs at (*t*^*^, *x*^*^). This gives the following solution:

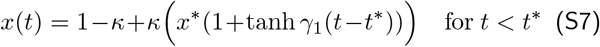

and

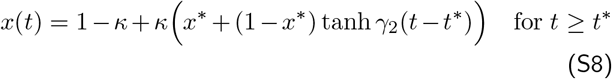

where

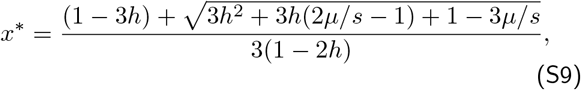

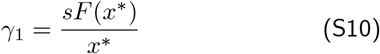

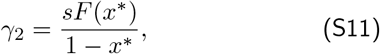

and *κ* relates to a correction so the solution matches the initial condition *x*(0) = *x*_0_:

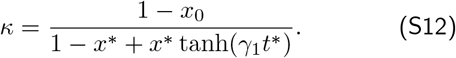

These can now be inserted into Eqn.S6 and integrated up to *t_K_*(*T*), to find the mean number of origins:

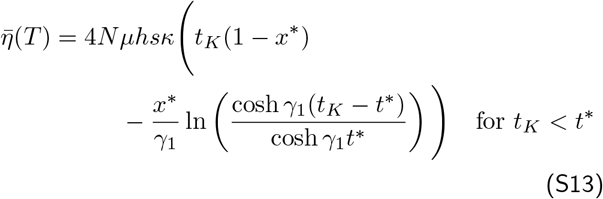

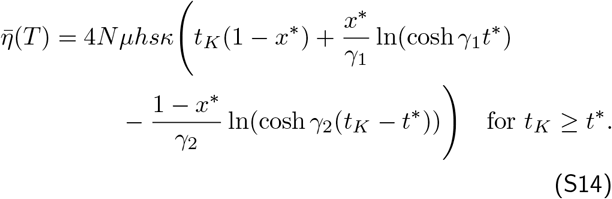

The likelihood in the diploid case is then given by the Poisson distribution:

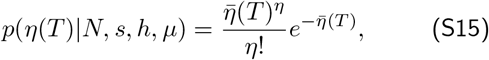

## References

[1] Anopheles gambiae 100 Genomes Consortium, 2017 Genetic diversity of the african malaria vector anopheles gambiae. Nature 552: 96.

[2] Bollback, J. P., T. L. York, and R. Nielsen, 2008 Estimation of 2nes from temporal allele frequency data. Genetics 179: 497–502.

[3] Bomblies, A., J.-B. Duchemin, and E. A. Eltahir, 2009 A mechanistic approach for accurate simulation of village scale malaria transmission. Malaria Journal 8: 223.

[4] Browning, S. and B. Browning, 2015 Accurate non-parametric estimation of recent effective population size from segments of identity by descent. The American Journal of Human Genetics 97: 404–418.

[5] Charlesworth, B., 2009 Effective population size and patterns of molecular evolution and variation. Nature Reviews Genetics 10: 195–205.

[6] Desai, M. M. and D. S. Fisher, 2007 Beneficial mutation selection balance and the effect of linkage on positive selection. Genetics 176: 1759–1798.

[7] Ewens, W. J., 2010 Mathematical Population Genetics: 1. A Theoretical Introduction. Springer.

[8] Feder, A. F., C. Kline, P. Polacino, M. Cottrell, A. D. M. Kashuba, B. F. Keele, S.-L. Hu, D. A. Petrov, P. S. Pen-nings, and Z. Ambrose, 2017 A spatio-temporal assessment of simian/human immunodeficiency virus (shiv) evolution reveals a highly dynamic process within the host. PLoS Pathogens.

[9] Fisher, R. A., 1930 The Genetical Theory of Natural Selection. Oxford Univ. Press, Oxford.

[10] Fu, X. L. Y.-X., 2015 Exploring population size changes using snp frequency spectra.

[11] Gutenkunst, R. N., R. Hernandez, S. Williamson, and C. Bustamante, 2009 Inferring the joint demographic history of multiple populations from multidimensional snp frequency data. PLoS Genetics.

[12] Hermisson, J. and P. S. Pennings, 2005 Soft sweeps: molecular population genetics of adaptation from standing genetic variation. Genetics 169: 2335–2352.

[13] Karasov, T., P. W. Messer, and D. A. Petrov, 2010 Evidence that adaptation in drosophila is not limited by mutation at single sites. PLoS genetics 6: e1000924.

[14] Keightley, P. D., R. W. Ness, D. L. Halligan, and P. R. Had-drill, 2014 Estimation of the spontaneous mutation rate per nucleotide site in a drosophila melanogaster full-sib family. Genetics 196: 313–320.

[15] Khatri, B. S., 2016 Quantifying evolutionary dynamics from variant-frequency time series. Scientific Reports 6.

[16] Kimura, M., 1962 On the probability of fixation of mutant genes in a population. Genetics 47: 713–719.

[17] Mabaso, M. L. H., T. Smith, A. Ross, and M. Craig, 2007 Environmental predictors of the seasonality of malaria transmission in africa: The challenge. The American Journal of Tropical Medicine and Hygiene 76: 33–38.

[18] Messer, P. W. and R. A. Neher, 2012 Estimating the strength of selective sweeps from deep population diversity data. Genetics 191: 593–605.

[19] Minakawa, N., G. Sonye, M. Mogi, A. Githeko, and G. Yan, 2002 The effects of climatic factors on the distribution and abundance of malaria vectors in kenya. Journal of Medical Entomology 51: 833–841.

[20] Pennings, P. S. and J. Hermisson, 2006a Soft sweeps ii-molecular population genetics of adaptation from recurrent mutation or migration. Molecular biology and evolution 23: 1076–1084.

[21] Pennings, P. S. and J. Hermisson, 2006b Soft sweeps iii: the signature of positive selection from recurrent mutation. PLoS genetics 2: e186.

[22] Petkova, D., J. Novembre, and M. Stephens, 2015 Visualizing spatial population structure with estimated effective migration surfaces. Nature Genetics 48: 94–100.

[23] Pybus, O. G., A. Rambaut, and P. H. Harvey, 2000 An integrated framework for the inference of viral population history from reconstructed genealogies. Genetics 155: 1429–37.

[24] Ralph, P. L. and G. Coop, 2010 Parallel adaptation: one or many waves of advance of an advantageous allele? Genetics

[25] Walker, M., P. Winskill, M.-G. B. nez, J. M. Mwangangi, C. Mbogo, J. C. Beier, and J. T. Midega, 2013 Temporal and micro-spatial heterogeneity in the distribution of anopheles vectors of malaria along the kenyan coast. Parasites & Vectors 6: 311.

[26] Waples, R. S. and C. Do, 2010 Linkage disequilibrium estimates of contemporary ne using highly variable genetic markers: a largely untapped resource for applied conservation and evolution. Evolutionary Applications 3: 244–262.

[27] Wilson, B. A., D. A. Petrov, and P. W. Messer, 2014 Soft selective sweeps in complex demographic scenarios. Genetics 198: 669–684.

[28] Wolfram Research, Inc., 2018 Mathematica, Version 11.3. Champaign, IL, 2018.

[29] Wright, S., 1931 Evolution in mendelian populations. Genetics 16: 97–159.

[30] Zanini, F., J. Brodin, L. Thebo, C. Lanz, G. Bratt, J. Albert, and R. A. Neher, 2015 Population genomics of intrapatient hiv-1 evolution. Elife 4: e11282.

